# One-shot learning with spiking neural networks

**DOI:** 10.1101/2020.06.17.156513

**Authors:** Franz Scherr, Christoph Stöckl, Wolfgang Maass

**Affiliations:** Institute of Theoretical Computer Science, University of Technology Graz

**Author notes:** {, }.

## Abstract

Understanding how one-shot learning can be accomplished through synaptic plasticity in neural networks of the brain is a major open problem. We propose that approximations to *BPTT* in recurrent networks of spiking neurons (RSNNs) such as *e-prop* cannot achieve this because their local synaptic plasticity is gated by learning signals that are rather ad hoc from a biological perspective: Random projections of instantaneously arising losses at the network outputs, analogously as in Broadcast Alignment for feedforward networks. In contrast, synaptic plasticity is gated in the brain by learning signals such as dopamine, which are emitted by specialized brain areas, e.g. VTA. These brain areas have arguably been optimized by evolution to gate synaptic plasticity in such a way that fast learning of survival-relevant tasks is enabled. We found that a corresponding model architecture, where learning signals are emitted by a separate RSNN that is optimized to facilitate fast learning, enables one-shot learning via local synaptic plasticity in RSNNs for large families of learning tasks. The same learning approach also supports fast spike-based learning of posterior probabilities of potential input sources, thereby providing a new basis for probabilistic reasoning in RSNNs. Our new learning approach also solves an open problem in neuromorphic engineering, where on-chip one-shot learning capability is highly desirable for spike-based neuromorphic devices, but could so far not be achieved. Our method can easily be mapped into neuromorphic hardware, and thereby solves this problem.

## Introduction

The most powerful methods for training a recurrent neural network rely on gradient-based optimization of a loss function *E* to obtain a well-performing set of network parameters *W*. The canonical way to compute gradients 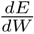 is to apply backpropagation through time (*BPTT*) (Werbos, 1990). However, it is widely believed that the brain does not employ *BPTT* for learning (Lillicrap and Santoro, 2019). Recently proposed alternatives to *BPTT*, such as *e-prop* (Bellec et al., 2019) for RSNNs and RFLO (Murray, 2019) for artificial neurons without slow processes focus on two types of variables that appear to be fundamental for the implementation of learning in the brain:

1. Neurons and synapses maintain traces of recent activity, which are known to induce synaptic plasticity if closely followed by a top-down learning signal (Yagishita et al., 2014; Gerstner et al., 2018). Such traces are commonly referred to as eligibility traces. We write 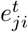 for the value of the eligibility trace for a synapse from neuron *i* to neuron *j* at time *t*.
2. There exists an abundance of top-down learning signals in the brain, both in the form of neuromodulators and in the form of firing activity (Sajad et al., 2019), some of them are known to be specific for different target populations of neurons, and to transmit a multitude of learning-relevant aspects (Roeper, 2013; Engelhard et al., 2019). We denote a learning signal to neuron *j* at time *t* by 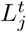.

The ideal weight change for network gradient descent in RSNNs can be expressed according to Bellec et al. (2019) as:

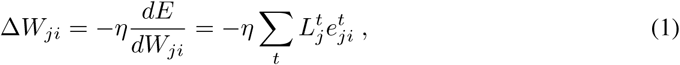

yielding the online synaptic plasticity rule 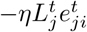 for time step *t*. The eligibility trace 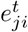 (more details are given later) is independent of the loss function *E* and just depends on the history of activations of the pre- and postsynaptic neuron. The ideal value of the learning signals 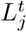 for implementing gradient descent is the total derivative 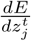, where 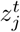 is a binary variable that assumes value 1 if neuron *j* spikes at time *t*, and otherwise value 0. But this learning signal is in general not available at time *t*, since it depends in general also on the impact of a spike at time *t* on values of the loss function *E* via future neuron outputs. In *random e-prop* it is replaced by a random projection of currently arising losses at the network outputs, like in Broadcast Alignment for feedforward networks (Lillicrap et al., 2016; Nøkland, 2016). This approximation to *BPTT* generally works quite well, but like *BPTT*, it typically requires a large number of training examples. We propose a more brain-like generation of learning signals for *e-prop*, where a separate, optimized RSNN emits learning signals. We found that this more natural setting, which we call *natural e-prop*, does in fact enable one-shot learning. We also found that it enables spike-based learning of posterior probabilities, thereby providing a new foundation for modeling brain-like probabilistic computations in RSNNs.

### Architectures

Our aim is to provide learning paradigms that can in principle be implemented on detailed models for neural networks of the brain. We focus here on simple standard models for spiking neurons, more precisely LIF neurons and a variation with Spike-Frequency-Adaptation (SFA) that we call ALIF neurons. The hidden variable of a LIF neuron is its membrane potential, which is a weighted sum –whose weights are the synaptic weights– of low-pass filtered versions of spike trains from presynaptic neurons. ALIF neurons (equivalent to the GLIF_2_ model of Teeter et al. (2018); Allen Institute: Cell Types Database (2018)) have an adaptive firing threshold as a second hidden variable. It increases for each spike of the neuron, and decays back to baseline with a time constant that is typically in the range of seconds according to biological data (Pozzorini et al., 2013, 2015). We included ALIF neurons in the RSNNs for Fig. 2 and 3 both because the presence of these neurons enhances temporal computing capabilities of RSNNs (Bellec et al., 2018; Salaj et al., 2020), and because a fraction of 20-40% of pyramidal cells in the neocortex exhibit SFA according to the database of the Allen Institute (Allen Institute: Cell Types Database, 2018). We used for simplicity fully connected RSNNs, but similar results can be achieved with more sparsely connected networks. We used these RSNNs to model both the Learning Network (LN) and the Learning Signal Generator (LSG) of our learning architecture, see Fig. 1A. External inputs to an RSNN were integrated into the membrane potential of neurons using weighted sums. The outputs (“readouts”) from the LN consisted of weighted sums of low-pass filtered spike trains from the neurons in the LN, modeling the impact of spikes in the LN on the membrane potential of downstream neurons in a qualitative manner. Their time constants are referred to as readout-time constants, and their impact on the readout value at an arbitrarily chosen time point *t* is indicated by the yellow shading in Fig. 1D, 2C, 3C. Importantly, these time constants were chosen to be relatively short to force the network to carry out spike based –rather than rate-based– computations. Simulations were carried out with a step size of 1 ms. Full details are provided in the Suppl.

**Figure 1:**
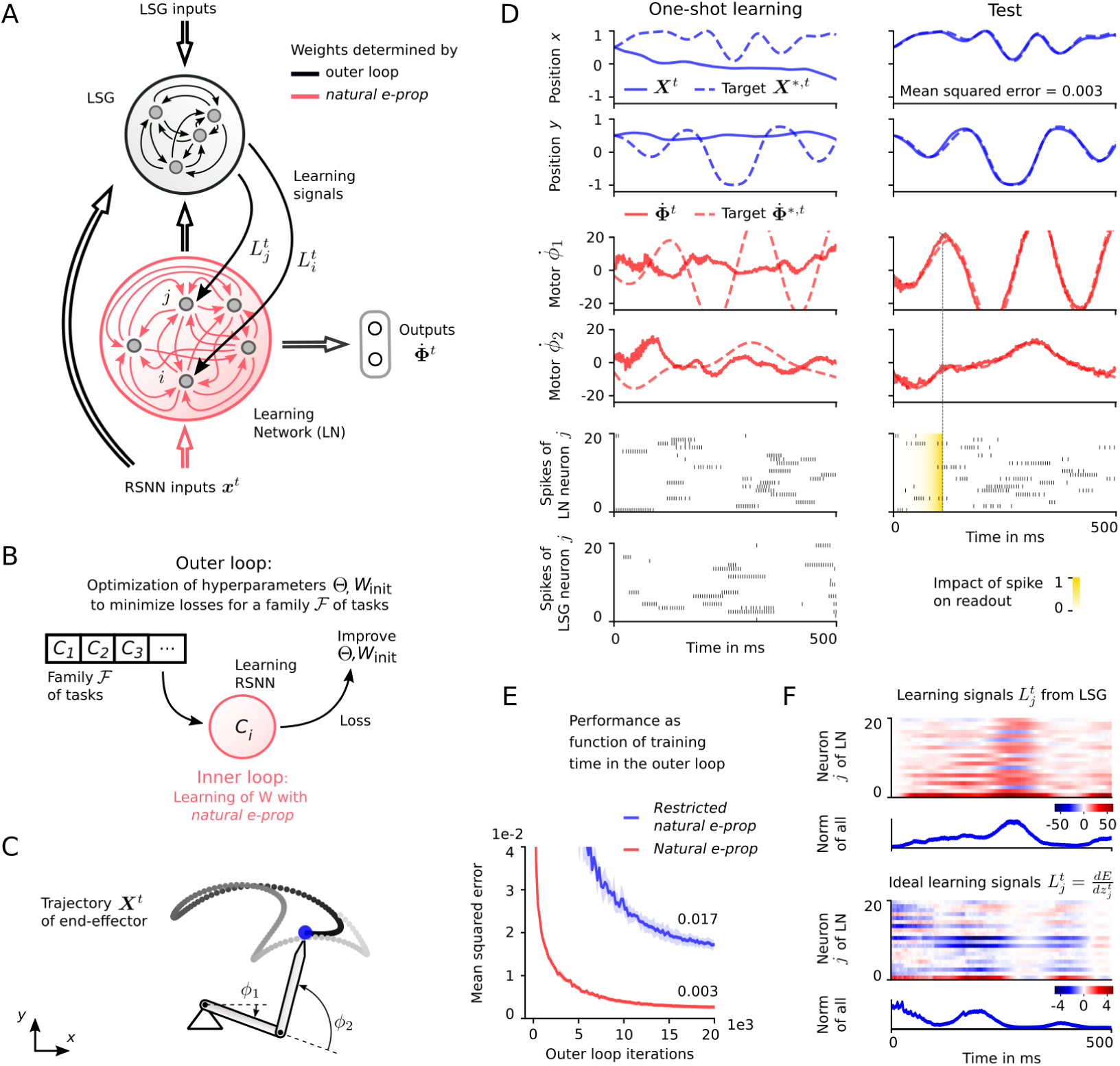
One-shot learning of novel arm movements with *natural e-prop*. **A)** Generic architecture for *natural e-prop*. The LN inputs ***x***^*t*^ consist in this case of a clock-like signal, and the outputs ***y***^*t*^ are velocities 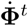 for the two joints of the arm. The LSG receives in addition to a copy of ***x***^*t*^ also the activity ***z***^*t*^ of the LN and the target movement ***X***^*,*t*^ as online network input. **B**) L2L scheme from Hochreiter et al. (2001), but used here with synaptic plasticity in the inner loop. **C**) Sample of a movement ***X***^*t*^, *t* = 1, …, 500 ms, of the end-effector of the arm. **D**) Demonstration of one-shot learning of a new arm movement ***X***^*,*t*^, *t* = 1, …, 500 ms: The single learning trial is shown on the left. The synaptic weights of the LN were updated according to *natural e-prop* after this single trial. The arm movement ***X***^*t*^ produced by the LN after this weight update is shown in the right column. Spike raster plots show a subset of neurons. **E**) MSE between target movement ***X***^*,*t*^ and ***X***^*t*^ in the testing trial shown on the right in D as function of the training time in the outer loop (mean and standard deviation (STD) obtained using 4 runs with different initial weights). **F**) The optimized LSG generates learning signals that differ substantially from the “ideal learning signals” used by *BPTT* under knowledge of the kinematic model.

### Algorithms

Learning by *natural e-prop* proceeded in two stages. During the first stage, corresponding to prior evolutionary and developmental optimization and to some extent also prior learning, we applied the standard learning-to-learn (L2L) or meta-learning paradigm (Fig. 1B). One considers there a very large –typically infinitely large– family ℱ of possibly relevant learning tasks *C*. Learning of a particular task *C* from ℱ by the LN was carried out in the inner loop of L2L via eqn (1), with the learning signals 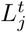 provided by the LSG. It was embedded during this first stage into an outer loop, where synaptic weights Θ related to the LSG (see black arcs in Fig. 1A) as well as the initial weights *W*_init_ of the LN were optimized via *BPTT*.

Specifically, every time the LN is first confronted with a new task *C* from the family ℱ, it starts with synaptic weights *W*_init_ and updates these according to (1). The learning performance of the LN on task *C* is evaluated with some loss function *E*_*C*_ (***y***^1^, …, ***y***^*T*^), where ***y***^*t*^ denotes the output from the LN at time *t* and *T* is the total duration that the network spends on task *C*. We then minimize the loss *E*_*C*_ over many task instances *C* that are randomly drawn from the family ℱ. This optimization in the outer loop was implemented through *BPTT*, like in (Bellec et al., 2018). In addition to the loss function described in the specific applications, we used regularization terms with the goal to bring the RSNNs into a sparse firing regime. Previous L2L applications used for learning in the inner loop either backpropagation (Andrychowicz et al., 2016; Finn et al., 2017), or no synaptic plasticity at all (Hochreiter et al., 2001; Wang et al., 2018; Bellec et al., 2018).

After this first stage of learning, all parameters regulated by the outer loop were kept fixed, and the learning performance of the LN was evaluated for new tasks *C* that were randomly drawn from ℱ. This performance is evaluated in Fig. 1E, 2D, 3D as function of the number of preceding iterations of the outer loop during stage 1.

Besides *natural e-prop*, we also tested the performance of a simplified version, called *restricted natural e-prop*. Like *random e-prop*, it uses no LSG and learning signals are weighted sums of instantaneously arising losses at the LN output. But in contrast to *random e-prop*, the weights of these error broadcasts –as well as the initial weights of the LN– are not chosen randomly but optimized in an outer loop of L2L that corresponds to the outer loop of *natural e-prop*.

The eligibility trace 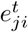 in (1) reflects the impact of the weight *W*_*ji*_ on the firing of the neuron *j* at time *t*, but only needs to take dependencies into account that do not involve other neurons besides *i* and *j*. For the case of a standard LIF neuron it is simply the product of a low-pass filtered version of the spike train from the presynaptic neuron *i* up to time *t* − 1 and a term that depends on the depolarization of the membrane of postsynaptic neuron *j* at time *t* (pseudo-derivative). For ALIF neurons the eligibility trace becomes a bit more complex, because it involves then also the temporal evolution of the use-dependent firing threshold of the neuron. More precisely, if 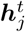 denotes the internal state vector of neuron *j*, i.e. the membrane voltage in the case of LIF neurons and additionally the dynamic state of the firing threshold in ALIF neurons, the eligibility trace is defined as 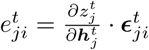. The quantity 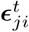 is a so-called eligibility vector and is recursively defined as: 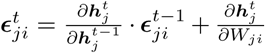, hence propagating an “eligibility” forward in time according to the dynamics of the postsynaptic neuron. Note that 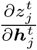 is in general not defined for a spiking neuron, and was therefore replaced by a “pseudo-derivative”, similar as in (Bellec et al., 2018). According to (Bellec et al., 2019) these normative eligibility traces are qualitatively similar to those that have been experimentally observed (Yagishita et al., 2014) and used in previous models for synaptic plasticity (Gerstner et al., 2018). They can be approximated by a fading memory of preceding pre-before-post firing events. Full details of the learning algorithms, regularization, as well as the values of hyperparameters that were used are given in the Suppl.

## Application 1: One-shot learning of new arm movements

Brea and Gerstner (2016) argue that one-shot learning is one of two really important learning capabilities of the brain that are not yet satisfactorily explained by current models in computational neuroscience. We demonstrate that *natural e-prop* supports one-shot learning in two very different contexts. We first consider a generic motor-control task, where the end-effector of a 2-joint arm has to trace a target trajectory that is given in Euclidean coordinates (*x, y*), see Fig. 1C. We show in Fig. 1D and E that a RSNN can learn this task with *natural e-prop* in one shot.

In order to perform the optimization of the LSG in the outer loop of L2L, we considered an entire family ℱ of tasks *C*. In each task, the RSNN was required to learn to reproduce a particular randomly generated target movement ***X***^*,*t*^ with the actual movement ***X***^*t*^ of the end-effector of the arm. The learning task was divided into two trials, a training and a testing trial, both starting out from the same initial state. During the training trial, the LSG was shown the target movement ***X***^*,*t*^ in Euclidean coordinates, and the LSG computed learning signals 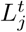 for the LN on the basis of ***X***^*,*t*^ and the spiking activity of the LN. After this trial, the accumulated weight update according to (1) was applied to the synapses of the LN. In the subsequent testing trial, one tested whether the LN was able to reproduce the previously demonstrated target movement of the end-effector of the arm –without receiving again the target trajectory ***X***^*,*t*^– while synaptic plasticity was turned off.

### Implementation

Feasible target movements ***X***^*,*t*^ of duration 500 ms were produced by application of randomly sampled target angular velocities 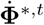 (sum of random sines) to the kinematic arm model. Each link of the arm had a length of 0.5. The LN consisted of 400 LIF neurons. Its input ***x***^*t*^ was the same across all trials and was given by a clock-like input signal: Every 100 ms another set of 2 input neurons started firing at 100 Hz. The output of the LN (with a readout-time constant of 20 ms) produced motor commands in the form of angular velocities of the joints 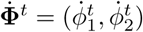. The LSG consisted of 300 LIF neurons. The online input to the LSG consisted of a copy of ***x***^*t*^, the activity ***z***^*t*^ of the LN, and the target movement ***X***^*,*t*^ in Euclidean coordinates. The LSG had no access to the errors resulting from the produced motor commands. For outer loop optimization we applied *BPTT* to 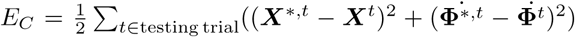. Note that ***X***^*t*^ was differentiable in our kinematic model. To compare with *BPTT* in Fig. 1F, we computed the “ideal learning signals” as 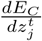 with *BPTT* over errors of the training trial. For *restricted natural e-prop* (see Fig. 1E) we defined the “instantaneously arising errors at network outputs” as the difference ***X***^*,*t*^ − ***X***^*t*^ in Euclidean coordinates. See Suppl. for full implementation details.

**Figure 2:**
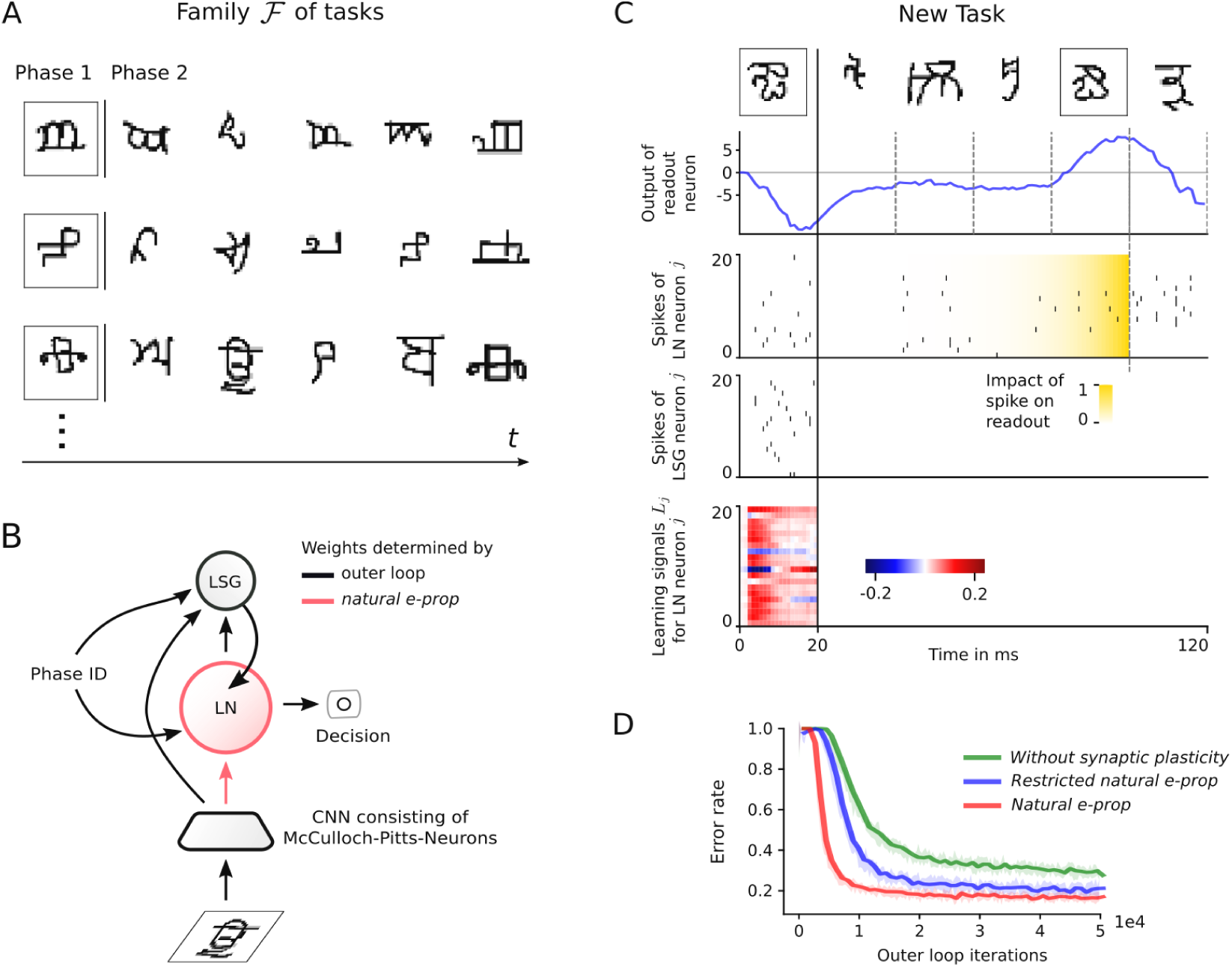
Biologically plausible model for learning a new class of characters from a single sample. **A)** Network inputs for three typical trials, with a single sample of a character class shown during phase 1, and samples from 5 character classes –including one from the same class as the sample during phase 1– shown during phase 2. For testing we used new character classes for all these samples, of which no sample had occurred during training in the outer loop. **B**) Input and output streams of the generic learning architecture for *natural e-prop* from Fig. 1A for this task. **C**) A sample trial for one-shot learning with *natural e-prop*. **D**) Error rates of *natural e-prop, restricted natural e-prop*, and an application of standard L2L without synaptic plasticity in the inner loop, as function of the duration of training in the outer loop (where the parameters of network components that are drawn black in B) are determined). Curve mean and STD were obtained using 10 runs with different initial weights.

**Figure 3:**
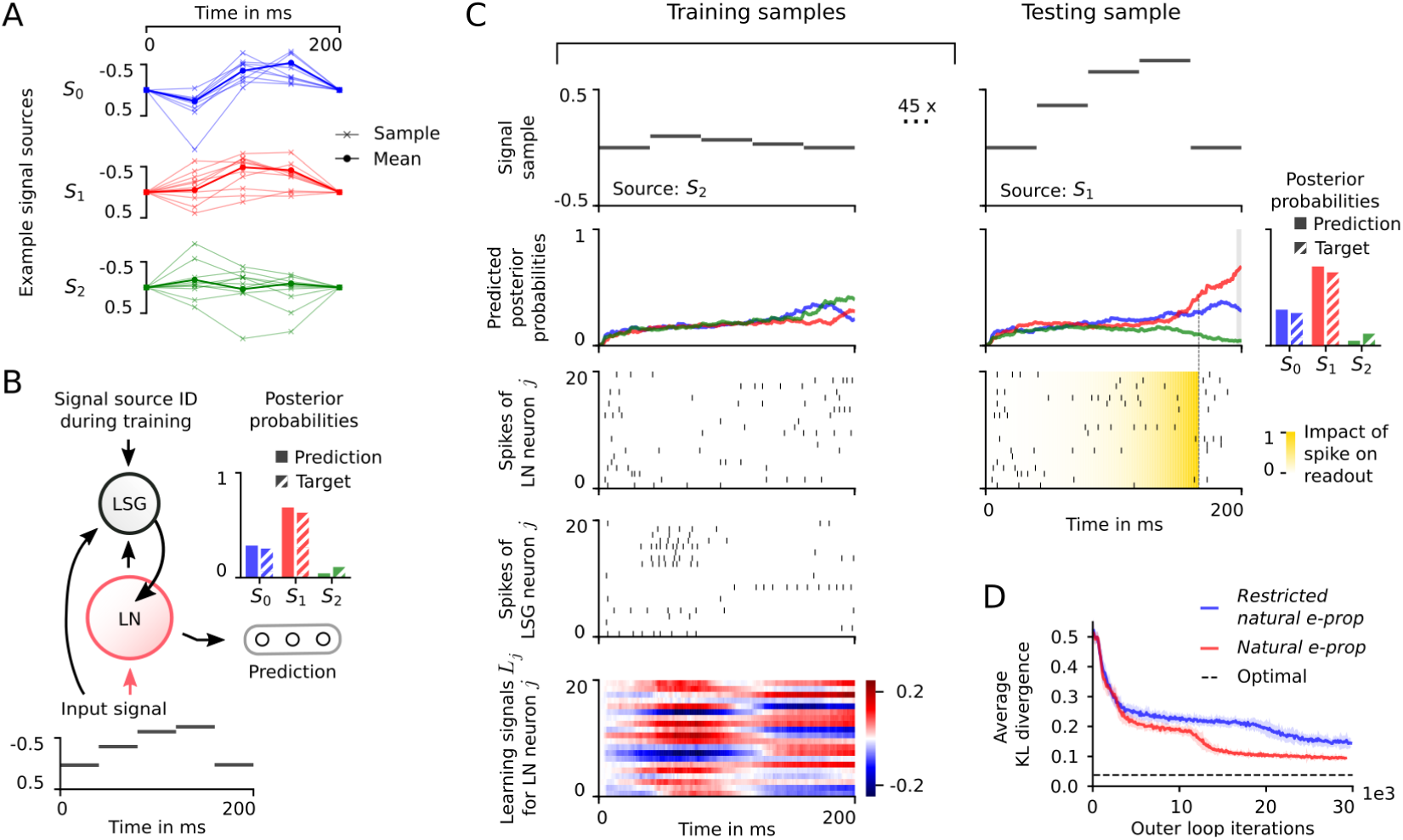
Fast spike-based learning of posterior probabilities with *natural e-prop*. **A)** Each task presented 3 new signal sources modeled as Gaussian processes (shown as interpolating lines for ease of interpretability). **B**) Architecture. **C**) Demonstration of fast learning of posterior probabilities: 1 out of 45 supervised signal samples is shown in the left column. 1 out of 5 test samples is shown in the right column. **D**) Average Kullback-Leibler (KL) divergence between target and predicted posterior probability distributions decreased during the course of outer loop optimization. LN predictions were normalized for computing KL divergence. Curve mean and STD were obtained using 32 runs with different initial weights. Optimal denotes the lower bound on KL divergence that can be achieved if one computes posterior probabilities in a Bayesian way, given all observed samples.

### Results

We tested the performance of *natural e-prop* for a random target movement (dashed curves in Fig. 1D), and show the trajectory ***X***^*t*^ produced by the LN as solid lines during the training and testing trial in the left and right column respectively. After the LSG had sent learning signals to the LN during the training trial and the resulting weight update in the LN, the LN could accurately reproduce the target movement using a biological realistic sparse firing activity (< 20 Hz).

Fig. 1E summarizes the mean squared error between the target ***X***^*,*t*^ and actual movement ***X***^*t*^ in the testing trial. The error is reported at different iterations of the outer loop optimization and decreases as the initial weights *W*_init_ of the RSNN and the weights Θ of the LSG become more adapted to the task family ℱ. Fig. 1E shows that *natural e-prop* accomplishes this one-shot learning task almost perfectly after a sufficiently good optimization of the LSG and initial weights of LN in the outer loop. It also shows that *restricted natural e-prop* works quite well, although with less precision. Fig. 1F shows that the learning signals that were emitted by the optimized LSG differed strongly from the “ideal learning signals” 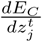 of *BPTT*. Hence, rather than approximating *BPTT*, the LSG appears to exploit common structure of a constrained range of learning tasks. This strategy endows *natural e-prop* with the potential to supersede *BPTT* for a concrete family ℱ of learning tasks.

Also a more general lesson can be extracted from the resulting solution of the motor control tasks: Whereas one usually assumes in models of biological motor control that forward and/or inverse models of the motor plant are indispensable, these are not needed in our model. Rather, the LSG finds during its evolution-like optimization a way to enable learning of precise motor control without these specialized modules. Hence our model provides a new perspective of the possible division of labor among subsystems of the brain that are involved in motor control and motor-related learning.

## Application 2: Learning a new class of characters from a single sample

The first of two challenges for building more human-like machines discussed in Lake et al. (2017) is the “characters challenge”, based on the well-known Omniglot dataset. It consists of 20 samples each of 1623 different classes of handwritten characters from 50 different alphabets. Each class of characters is seen in a more general context as a prototypical visual concept, such as a new type of vehicle or the face of a new person. For one-shot learning, a single sample of a new class is presented during phase 1. After this sample disappears, the learner gets to see in phase 2 samples from the same and other new classes of characters, and has to pick the one which is from the same class as the one shown in phase 1 (see Fig. 2A). In the simpler version of this task that is considered in Fig. 1A in Lake et al. (2017, 2019) all samples are presented during phase 2 simultaneously, and the learner has to solve a multiple-choice test. In other words, the learner can view and compare the samples repeatedly and in any order. We considered a more difficult and arguably more biological online version of this one-shot learning task, where the samples are presented during phase 2 sequentially, including exactly one from the same class as the sample from phase 1. The learner has to decide for each sample instantaneously whether it belongs to the same class as the sample from phase 1. The task is solved correctly only if this decision is correct for all 5 samples shown during phase 2, otherwise the performance is counted as error. Informal testing of our human performance on this task yielded error rates around 15% (based on 4 subjects, each carrying out 100 trials; code for testing one’s own performance in Suppl.). Numerous methods for solving the simpler version of this task are discussed in the review (Lake et al., 2019). Methods based on probabilistic program induction (Lake et al., 2015) and variants of artificial neural networks such as Matching Networks (Vinyals et al., 2016) and Prototypical Networks (Snell et al., 2017) perform especially well. We are not aware of any biologically plausible neural network architecture or learning method that can solve this task.

### Implementation

The LN consisted of 266 LIF and 181 ALIF neurons. Its input ***x***^*t*^ consisted of the output of a 3-layer CNN consisting of 15488 binary neurons (i.e., McCulloch-Pitts neurons or threshold gates chosen here instead of spiking neurons to save compute time), whose weights are trained in the outer loop of L2L (see Fig. 2B). Note that we believe that the binary neurons in the CNN could be made spiking with enough compute resources. The input ***x***^*t*^ also included the phase ID, which was binary encoded with value 0 for phase 1, switching to a value of 1 for phase 2. The characters from the Omniglot dataset were presented to the CNN in the form of 28 x 28 arrays of grayscale pixels. A single output (with a readout-time constant of 10 ms) was used for deciding online during phase 2 whether the just seen character belonged to the same class as the one from phase 1. Its analog output value at the time points indicated by dashed lines (see 2nd row of Fig. 2C) was taken as decision that the just seen character was from the same class as the one from phase 1 if it had a positive value. Each character –including the one in phase 1– was presented for just 20 ms, thereby forcing the LN to carry out spike-based rather than rate-based-computation and learning. Another more practical advantage of these short presentation times were savings in the compute time needed for the experiment. The LSG consisted of 149 LIF and 100 ALIF neurons. The input to the LSG entailed a copy of ***x***^*t*^ (which included the phase ID) and the activity ***z***^*t*^ of the LN, The LSG sent learning signals to the LN only during the first phase and the accumulated weight update according to (1) was realized at the end of phase 1. For optimization in the outer loop, *BPTT* was applied on the cross-entropy loss.

### Results

A sample trial for one-shot learning of a new class of characters via *natural e-prop* is shown in Fig. 2C. Synaptic plasticity according to (1) takes place just once, at the end of phase 1 of the trial (the activity of the LSG during phase 2 does not matter, hence we simply turned it off during phase 2). Note that the yellow shading Fig 2 makes clear that also makes clear that the decision results from a spike-based rather than rate-based regime, especially in view of the sparse firing rate of the LN of 20 Hz.

The resulting error rates for new tasks (with character classes that did not occur during training in the outer loop) is shown in Fig. 2D. *Natural e-prop* reaches after sufficiently long training in the outer loop an error rate of 16.2%, which is close to our estimate of human error rates around 15%. Restricted *natural e-prop* produced a higher error rate of 19.7%. For comparison we also plotted the performance achieved by the standard version of L2L without synaptic plasticity of the LN in the inner loop. It achieved a considerable higher error rate of 28.4% Altogether this result shows that a biologically plausible learning method, *natural e-prop*, applied to relatively small generic RSNNs can solve this one-shot learning task at a close-to-human performance level.

## Application 3: Fast learning of posterior probabilities

The ability of neural circuits in the brain to represent and manipulate probability distributions lies at the heart of numerous models of brain computation (Rao et al., 2002; Doya et al., 2007). We show here that RSNNs can learn with *natural e-prop* to estimate and represent such probabilities on the fast level of spikes, rather than firing rates. We demonstrate this for a task that arguably is among the most salient ones for generic neural circuits in virtually any brain area, that can rarely be certain about the sources of signals which they are currently receiving: To estimate the posterior probabilities of different potential causes of time-dependent input signals *U*. Estimation of categorical probabilities with raw network outputs is a challenging task for a RSNN if only samples of such a distribution are accessible, because the cross-entropy loss does not apply to unnormalized values. With *natural e-prop* on the other hand, the LSG can acquire the capability to produce only such learning signals that let the network predict suitably normalized probability values.

### Task

We chose a family ℱ of tasks *C*, each defined by 3 new signal sources *S*_*i*_ with random characteristics: Gaussian process generators with a time-varying mean, see Fig. 3A for an example set of signal sources. The task of the LN was to assign posterior probabilities *p*(*S*_*i*_ | *U*) to signal samples *U* from a randomly selected source *S*_*i*_, which characterized how likely a source *S*_*i*_ had generated the signal *U* (Fig. 3B). The LN had to produce these probabilities after having received just 45 sample signals with known source *S*_*i*_.

### Implementation

Signals *U* = (*u*^1^, …, *u*^200^) had a duration of 200 ms, where *u*^*t*^ was constant for consecutive 40 ms. The resulting 5 components were samples of a Gaussians distribution using a covariance matrix generated by a rational quadratic kernel (Gaussian process). The first and last component were conditioned to be 0. The value *u*^*t*^ of the signal was encoded by 100 analog values and given as input ***x***^*t*^ to the LN (see Suppl.). The input ***x***^*t*^ also included the binary value that switched from 0 to 1 after 120 ms. The LN consisted of 340 LIF and 160 ALIF neurons. It produced predictions of posterior probabilities via 3 outputs with a readout-time constant of ∼30 ms whose values were taken as predictions after 200 ms, using no normalization. The LSG consisted of 180 LIF neurons and 120 ALIF neurons. It received during supervised samples a copy of ***x***^*t*^, the activity in the LN ***z***^*t*^ and the signal source identity as inputs (Fig. 3B). After the 45 supervised samples, the weight update according to (1) was applied, and the LN was tested on 5 more signal samples. The LSG was inhibited during this testing phase. The loss *E*_*C*_ for optimization in the outer loop was based on the cross-entropy between target and predicted posterior probabilities on the 5 testing samples. Additional penalties ensured that the LN produced valid probability distributions. In the case of *restricted natural e-prop* learning signals existed only at the last time step at *t* = 200 ms, and were defined as a weighted broadcast of the difference between the prediction and the one-hot encoded signal source identity. See Suppl. for all details of the implementation.

### Results

Fig. 3D shows that *natural e-prop* enables RSNNs to estimate posterior probabilities for possible sources of input time series quite fast-on the level of spikes, operating in a sparse firing regime with rates below 20 Hz - and in a rather sample efficient manner (see Bayesian optimum as baseline). Intriguingly, no explicit computation of softmax or any form of normalization was required for the RSNN. Rather, an LSG can acquire through evolution-like optimization the capability to induce RSNNs to learn good estimates of posterior probabilities without that. The LN produced outputs *y*_*i*_ that were normalized with high precision (at time *t* = 200 ms): The average error on normalization (Σ _*i*_ *y*_*i*_ − 1)^2^ was (8 *±* 10) *·* 10^−4^.

## Discussion

We found that if one combines local rules for synaptic plasticity with learning signals from a separate network that is optimized for inducing fast learning of RSNNs, one-shot learning becomes feasible in a biologically realistic manner. This solves a well-known open problem in computational neuroscience (Brea and Gerstner, 2016). We have also shown that RSNNs can learn in this way to estimate posterior probabilities, thereby providing a new basis for modeling probabilistic computing and learning in RSNNs. A while ago it was commonly assumed that learning signals that are provided by dopaminergic neurons in the brain, e.g. from VTA, send uniform learning signals with a single message –reward prediction error– to diverse neural networks of the brain. But recent experimental data show that these learning signals are substantially more complex and diverse (Engelhard et al., 2019). Our approach suggests that these learning signals from VTA and other specialized brain areas have been optimized by evolution to enable other brain areas to exhibit diverse advanced learning capabilities, such as one-shot learning and efficient estimation of probabilities. Incidentally, these specialized learning capabilities are also desirable for neuromorphic hardware, and our approach opens the door for implementing one-shot learning in spike-based neuromorphic hardware.

## Acknowledgments

This research/project was supported by the Human Brain Project (Grant Agreement number 785907) of the European Union and a grant from Intel. We gratefully acknowledge the support of NVIDIA Corporation with the donation of the Quadro P6000 GPU used for this research. This work was supported by a grant from the Swiss National Supercomputing Centre (CSCS) under project ID ich024. We would like to thank Sandra Diaz from the SimLab at the FZ Jülich for enabling the use of CSCS. We thank Martin Pernull, Philipp Plank and Anand Subramoney for helpful comments on an earlier version of the manuscript. Special thanks go to Guillaume Bellec for his insightful comments and ideas when carrying out this work.

## Notes

### Competing Interest Statement

The authors have declared no competing interest.

