## Supplementary text, figures and tables for "One-shot learning with spiking neural networks"

### Supplements for “One-shot learning with spiking neural networks”

**Franz Scherr, Christoph Stöckl and Wolfgang Maass**
Institute of Theoretical Computer Science
University of Technology Graz
{scherr,stoeckl,maass}@igi.tugraz.at

**S3 Computing Infrastructure** **4**

**S4 Details of application 1: one-shot learning of new arm movements** **5**

**S5 Details of application 2: learning a new class of characters from a single sample** **5**

**S6 Details of application 3: learning of posterior probabilities** **8**

Preprint. Under review.

#### S1 Architectures

##### S1.1 RSNNs

We modeled the RSNNs in our experiments using standard LIF neurons and a variation with Spike-Frequency-Adaptation (SFA) that we call ALIF neurons. Outputs from the LN and outputs from the LSG (learning signals) are modeled by weighted low-pass filtered network spikes. We considered densely connected RSNNs, meaning that every input is connected to every neuron in the RSNN, and the neurons in the RSNN are themselves connected all-to-all. We did not consider that a neuron is connected to itself by a synapse (autapse).

**LIF** In this formulation, every neuron $j$ is equipped with a hidden variable: its membrane voltage $v_j^t$ at time $t$. This membrane voltage integrates input currents and decays back to a resting potential according to its membrane time constant $\tau_m$. Every time $v_j^t$ crosses the threshold voltage $v_{\text{th}}$ from below, the neuron produces an output $z_j^t = 1$, called a spike, and remains silent otherwise. In discrete time steps of size $\delta t$ ($= 1$ ms in our simulations), a LIF neuron is characterized by the equations:

$$v_j^t = \alpha v_j^{t-1} + \sum_i W_{ji}^{\text{in}} x_i^t + \sum_{i \neq j} W_{ji}^{\text{rec}} z_i^{t-d} - z_j^{t-1} v_{\text{th}} \ , \quad \text{(S1)}$$

$$z_j^t = H(v_j^t - v_{\text{th}}) \ . \quad \text{(S2)}$$

Here $W_{ji}^{\text{rec}}$ is the synaptic weight from network neuron $i$ to $j$ and $W_{ji}^{\text{in}}$ is the weight of input component $x_i^t$ for neuron $j$, the factor $\alpha = e^{-\delta t/\tau_m}$ describes how quickly the membrane voltage decays, $H$ denotes the Heaviside step function and $d$ is the transmission delay of recurrent spikes (in the range of 1 to 5 ms in our experiments). We also used a simple model for a refractory period, where we set $z_j^t = 0$ after a spike of neuron $j$ for a time $t_{\text{refrac}}$ (in the range of 2 to 5 ms in our experiments).

**ALIF** ALIF neurons have a second hidden variable besides the membrane potential $v_j^t$ that accounts for the variable component of the firing threshold and is denoted by $a_j^t$. In this case, the resulting threshold voltage $A_j^t$ increases with every output spike and decays back to the baseline threshold $v_{\text{th}}$ according to an adaptation time constant $\tau_a$ (typically on the order of 200 ms in our simulations). Hence, for ALIF neurons, equ. (S2) is replaced by:

$$z_j^t = H(v_j^t - A_j^t) \ , \quad \text{(S3)}$$

$$A_j^t = \beta a_j^t + v_{\text{th}} \ , \quad \text{(S4)}$$

$$a_j^t = \rho a_j^{t-1} + z_j^{t-1} \ , \quad \text{(S5)}$$

where $\rho = e^{-\delta t/\tau_a}$. The factor $\beta$ denotes the impact of threshold adaptation.

**Output** Outputs from the LN and from the LSG are constructed by a weighted sum of low-pass filtered network spikes. LN outputs are defined by:

$$y_k^t = (1 - \nu) \sum_{t' \leq t} \sum_j \nu^{t-t'} W_{kj}^{\text{out}} z_j^{t'} + b_k^{\text{out}} \ , \quad \text{(S6)}$$

with output weights $W_{kj}^{\text{out}}$, output biases $b_k^{\text{out}}$, $\nu = e^{-\delta t/\tau_{\text{out}}}$ and $\tau_{\text{out}}$ is the readout-time constant. Learning signals are defined similar but use spikes of the LSG, a different set of output weights and biases, and a different readout-time constant $\tau_{\text{learning signals}}$.

##### S1.2 Eligibility traces

Each synapse from neuron or input $i$ to $j$ has an associated eligibility trace, which is a core concept of *e-prop* (Bellec et al., 2019). The eligibility trace $e_{ji}^t$ reflects the impact of the weight $W_{ji}$ on the firing of the neuron $j$ at time $t$, but only needs to take dependencies into account that do not involve other neurons besides $i$ and $j$. More precisely, eligibility traces exist separately for input and recurrent weights. Let $\boldsymbol{h}_j^t$ denote the hidden variables for a neuron $j$ at time $t$, i.e. the membrane voltage in the case of LIF neurons and additionally the dynamic state of the firing threshold in ALIF neurons. The eligibility trace is then defined by:

$$e_{ji}^t = \frac{\partial z_j^t}{\partial \boldsymbol{h}_j^t} \cdot \boldsymbol{\epsilon}_{ji}^t \,, \tag{S7}$$

$$\boldsymbol{\epsilon}_{ji}^t = \frac{\partial \boldsymbol{h}_j^t}{\partial \boldsymbol{h}_j^{t-1}} \cdot \boldsymbol{\epsilon}_{ji}^{t-1} + \frac{\partial \boldsymbol{h}_j^t}{\partial W_{ji}} \,. \tag{S8}$$

The recursive definition of the so-called eligibility vector $\boldsymbol{\epsilon}_{ji}^t$ visualizes that this quantity can be propagated forward in time along with network computation. For RSNNs the term $\frac{\partial z_j^t}{\partial \boldsymbol{h}_j^t}$ is problematic because the relationship between $z_j^t$ and $\boldsymbol{h}_j^t$ includes the non-differentiable Heaviside function. We replace its derivative in equ. (S7) with a "pseudo-derivative" $\psi_j^t = 0.3 \cdot \max\left(0, 1 - \left|\frac{v_{\mathrm{th}} - v_j^t}{v_{\mathrm{th}}}\right|\right)$ (more details can be found in S2.1).

**Eligibility traces for postsynaptic LIF neurons** The vector of hidden variables $\boldsymbol{h}_j^t$ is for LIF neurons simply the membrane potential $\boldsymbol{h}_j^t = (v_j^t)$ and the application of the definition of eligibility traces in equ. (S7-S8) to the definition of LIF dynamics in equ. (S1-S2) yields:

$$e_{ji}^t = \psi_j^t \bar{z}_i^{t-d}, \tag{S9}$$

for a synapse of neuron $i$ to neuron $j$, where we additionally define $\bar{z}_i^t = \sum_{t' \leq t} \alpha^{t-t'} z_i^{t'}$ as the low-pass filtered presynaptic spike train of neuron $i$. For input weights, we simply replace $\bar{z}_i^{t-d}$ by $\bar{x}_i^t$.

**Eligibility traces for postsynaptic ALIF neurons** In this case, the vector of hidden variables of a neuron also includes the variable component of the firing threshold: $\boldsymbol{h}_j^t = (v_j^t, a_j^t)$. The derivative $\frac{\partial \boldsymbol{h}_j^t}{\partial \boldsymbol{h}_j^{t-1}}$ is now given by a 2x2 matrix. The first row is associated to the dynamics of the membrane voltage and its entries are $\frac{\partial v_j^t}{\partial v_j^{t-1}} = \alpha$ and $\frac{\partial v_j^t}{\partial a_j^{t-1}} = 0$. We proceed similarly for the dynamics of the threshold adaptation but replace the spike $z_j^{t-1}$ in equ. (S5) by its definition to capture the impact of the membrane voltage on threshold adaptation. Hence we find for the entries on the second row: $\frac{\partial a_j^t}{\partial a_j^{t-1}} = \rho - \beta \psi_j^{t-1}$ and $\frac{\partial a_j^t}{\partial v_j^{t-1}} = \psi_j^{t-1}$. This yields the following representation of the eligibility trace $e_{ji}^t$ for the connection from neuron $i$ to an ALIF neuron $j$:

$$e_{ji}^t = \psi_j^t (\bar{z}_i^{t-d} - \beta \epsilon_{a,ji}^t) \,, \tag{S10}$$

$$\epsilon_{a,ji}^t = (\rho - \beta \psi_j^{t-1}) \epsilon_{a,ji}^{t-1} + \psi_j^{t-1} \bar{z}_i^{t-d-1} \,. \tag{S11}$$

**Eligibility traces and refractory period** To account for the refractory period of neurons in eligibility traces, we simply set the "pseudo-derivative" to zero $\psi_j^t = 0$ at times $t$ when neuron $j$ is refractory.

### S2 Algorithms

#### S2.1 Details of outer loop optimization

Optimization on the level of the outer loop was implemented using gradient descent in RSNNs. More precisely, we considered that the entire inner loop, consisting of a task, an RSNN and of plasticity applied to the RSNN, is one large dynamical system for which we compute gradients using *BPTT*. To accomplish this, we replaced the derivative $\frac{\partial H(v_j^t - v_{\mathrm{th}})}{\partial v_j^t}$ of the Heaviside function in equ. (S2) with a "pseudo-derivative" in the backward pass according to Bellec et al. (2018), which was defined by:

$$\psi_j^t = 0.3 \cdot \max\left(0, 1 - \left|\frac{v_{\mathrm{th}} - v_j^t}{v_{\mathrm{th}}}\right|\right) \,. \tag{S12}$$

Similarly, for ALIF neurons, we replaced the derivative $\frac{\partial H(v_j^t - A_j^t)}{\partial v_j^t}$ of the Heaviside function in equ. (S3) by $0.3 \cdot \max\left(0, 1 - \left|\frac{A_j^t - v_j^t}{v_{\text{th}}}\right|\right)$. We simulated and computed gradients for a batch of $N_{\text{batch}}$ different instances of the inner loop and actual updates to the initial weights $W_{\text{init}}$ of the LN and weights $\Theta$ of the LSG were realized by application of Adam (Kingma and Ba, 2014) with a learning rate of $\eta_{\text{outer loop}}$.

##### S2.2 Regularization

In order to bring the RSNNs into a sparse firing regime, we added additional terms for regularization of activity to the outer loop loss $E_C$. We applied different types of regularization.

**Firing rate regularization** To keep the average firing rate $f_j$ for all neurons $j$ in the RSNN close to a predefined target firing rate $f_{\text{target}}$ (either 10 Hz or 20 Hz in our experiments), we added a term $\lambda_f E_{\text{rate}}$ defined by:

$$\lambda_f E_{\text{rate}} = \lambda_f \sum_j (f_j - f_{\text{target}})^2 \ . \tag{S13}$$

We computed $f_j$ as the average spike count across batch and time: $f_j = \frac{1}{N_{\text{batch}} T} \sum_{n=1}^{N_{\text{batch}}} \sum_{t=1}^{T} z_j^{(n,t)}$, where $z_j^{(n,t)}$ additionally indexes spikes of a neuron in a particular batch with $n$ (recall that $T$ is the total duration the LN spends on a particular task $C$). The factor $\lambda_f$ is a hyperparameter that scales the importance of firing rate regularization.

**Voltage range regularization** Similarly, in order to encourage the membrane voltage to remain in a particular range, we penalized values of the membrane voltage that leave this range. We added a term $\lambda_v E_v$ defined by:

$$\lambda_v E_v = \frac{\lambda_v}{N_{\text{batch}} T} \sum_{n=1}^{N_{\text{batch}}} \sum_{t=1}^{T} \sum_j \left( \max(0, v_j^{(n,t)} - A_j^{(n,t)})^2 + \max(0, -v_j^{(n,t)} - v_{\text{th}})^2 \right) \ . \tag{S14}$$

Note that we again used an index $n$ to access individual batches. For LIF neurons $A_j^{(n,t)}$ is replaced by $v_{\text{th}}$. The factor $\lambda_v$ is a hyperparameter that scales the importance of the resulting voltage range regularization.

##### S2.3 *Restricted natural e-prop*

Due to the readout-time constant $\tau_{\text{out}}$, a spike $z_j^t$ has a temporally extended impact on the output. For this reason, Bellec et al. (2019) derived that *random e-prop* requires in equ. (1) a low-pass filtered eligibility trace $\bar{e}_{ji}^t = \sum_{t' \leq t} \nu^{t-t'} e_{ji}^{t'}$ that takes into account the decay of the output when combined with weighted sums of instantaneously arising output errors. If not mentioned otherwise, we adopted the same strategy for *restricted random e-prop* and used low-pass filtered eligibility traces together with optimized weighted error broadcasts. This yields the weight update rule for *restricted natural e-prop*:

$$\Delta W_{ji} = -\eta \sum_t L_j^t \bar{e}_{ji}^t \ . \tag{S15}$$

We did not observe a benefit of low-pass filtered eligibility traces in the case of *natural e-prop*.

#### S3 Computing Infrastructure

Experiments were carried out using Tensorflow (Abadi et al., 2016). We used accelerated hardware in the form of GPUs of type Nvidia GeForce GTX 1080Ti and Nvidia Tesla P100.

#### S4 Details of application 1: one-shot learning of new arm movements

**Details of the task** In this task, the two outputs of the LN were interpreted as angular velocities $\dot{\mathbf{\Phi}}^t = (\dot{\phi}_1^t, \dot{\phi}_2^t)$, which were applied to the joints of the kinematic arm model. The configuration of the arm model at time $t$ is fully described by the angles of the joints $\mathbf{\Phi}^t = (\phi_1^t, \phi_2^t)$. These angles are measured against the horizontal and the first leg of the arm respectively, see Figure 1C. For given angles $\mathbf{\Phi}^t$, the Euclidean coordinates $\boldsymbol{X}^t = (x^t, y^t)$ of the tip of the arm are defined by $x^t = l\cos(\phi_1^t) + l\cos(\phi_1^t + \phi_2^t)$ and $y^t = l\sin(\phi_1^t) + l\sin(\phi_1^t + \phi_2^t)$ with $l = 0.5$. The kinematic arm model was simulated by Euler integration: $\phi_i^t = \sum_{t' \leq t} \dot{\phi}_i^{t'} \delta t + \phi_i^0$ using a $\delta t = 1\,\mathrm{ms}$. The initial values were set to $\phi_1^0 = 0$ and $\phi_2^0 = \frac{\pi}{2}$. Feasible target movements $\boldsymbol{X}^{*,t}$ of duration 500 ms were created randomly by sampling the generating angular velocities $\dot{\mathbf{\Phi}}^{*,t} = (\dot{\phi}_1^{*,t}, \dot{\phi}_2^{*,t})$. Each component was given by:

$$\dot{\phi}_i^{*,t} = \sum_m S_{im} \sin\left(2\pi\omega_{im}\frac{t}{T} + \delta_{im}\right) \stackrel{\text{def}}{=} \sum_m q_{im}^t \, , \tag{S16}$$

where $m$ was set to 5, $S_{im}$ was sampled uniformly in $[0, 30]$, $\omega_{im}$ was sampled uniformly in $[0.3, 1]$ and $\delta_{im}$ was sampled uniformly in $[0, 2\pi]$. After this sampling, every component $q_{2,m}^t$ in $\dot{\phi}_2^{*,t}$ was rescaled to satisfy $\max_t(q_{2,m}^t) - \min_t(q_{2,m}^t) = 20$. In addition, we considered constraints on the angles of the joints: $\phi_1 \in [-\frac{\pi}{2}, \frac{\pi}{2}]$ and $\phi_2 \in [0, \pi]$. If violated, the respective motor commands $\dot{\phi}_i^{*,t}$ were rescaled to match the constraints.

**Architecture** The LN consisted of $N_{\mathrm{LN}}$ neurons and the LSG consisted of $N_{\mathrm{LSG}}$ neurons. See Tab. S1 for the values used in each experiment.

**Input encoding** The input $\boldsymbol{x}^t$ to the LN was the same for each trial and was given by 10 input neurons that produced a clock-like input input signal: Every 100 ms another set of 2 input neurons started firing regular spike trains of 100 Hz.

The online input to the LSG consisted of a copy of $\boldsymbol{x}^t$, the activity $\boldsymbol{z}^t$ of the LN, and the target movement $\boldsymbol{X}^{*,t} = (x^{*,t}, y^{*,t})$ in Euclidean coordinates. The 2 Euclidean coordinates were encoded using a population of 100 neurons each, where each neuron in such a population fired for a preferred scalar value. More precisely, for encoding the coordinate $x^{*,t}$, a neuron $i \in \{1, \ldots, 100\}$ in the corresponding population emitted a Poisson spike train with an instantaneous rate of $\exp\left(-\frac{(x^{*,t} - (i \cdot \frac{2.8}{99} - 1.4))^2}{2 \cdot 0.2^2}\right) \cdot 200$ Hz. $y^{*,t}$ was encoded similarly.

***Restricted natural e-prop*** The learning signal in *restricted natural e-prop* for a neuron $j$ was defined by:

$$L_j^t = \sum_k B_{jk}(X_k^t - X_k^{*,t}) \, , \tag{S17}$$

with $B_{jk}$ being optimized in the outer loop. Note that the sum over $k$ is over the 2 coordinates $x, y$. We did not use low-pass filtered eligibility traces $\bar{e}_{ji}^t$ in this case.

**Details of outer loop optimization** In both the training and the testing trial, the LN started out from the same initial state, which was defined by $v_j^0 = 0$ for all neurons $j$. Additionally, the learning rate in the outer loop was reduced every 300 outer loop iterations to a fraction of 0.95 of the previous value. The outer loop optimization procedure approximately required 36 hours for $20 \cdot 10^3$ iterations on a single Nvidia GeForce GTX 1080Ti.

**Hyperparameters** Hyperparameters were tuned manually for this application and we display our setting and associated search range in Tab. S1.

#### S5 Details of application 2: learning a new class of characters from a single sample

**Details of the task** Each task $C$ consisted of a sequence of images where in phase 2 exactly one image of the same class as the one shown in phase 1 appeared. To construct such sequences, we

Table S1: List of hyperparameters used in application 1. Note that $N_{\text{batch}}$, $d$, $t_{\text{refrac}}$,$N_{\text{LN}}$ and $N_{\text{LSG}}$ are integer parameters.

*Natural e-prop*

| Symbol | Name | Value | Search range |
|---|---|---|---|
| $\tau_{\text{learning signals}}$ | Readout-time constant of learning signals | 20 ms | [10 ms, 30 ms] |
| $N_{\text{LSG}}$ | Number of neurons in the LSG | 300 | {300} |

Common hyperparameters

| Symbol | Name | Value | Search range |
|---|---|---|---|
| $\tau_m$ | Membrane time constant | 20 ms | $\{15\text{ ms}, 20\text{ ms}\}$ |
| $\tau_{\text{out}}$ | Readout-time constant | 20 ms | $\{15\text{ ms}, 20\text{ ms}\}$ |
| $d$ | Synaptic transmission delay | 1 ms | {1 ms} |
| $t_{\text{refrac}}$ | Duration of refractory period | 5 ms | $\{1\text{ ms}, 5\text{ ms}\}$ |
| $\eta$ | Learning rate in inner loop | $10^{-4}$ | [$10^{-4}$,$10^{-3}$] |
| $\eta_{\text{outer loop}}$ | Learning rate in outer loop | $1.5 \cdot 10^{-3}$ | $[1 \cdot 10^{-3}, 2 \cdot 10^{-3}]$ |
| $N_{\text{batch}}$ | Batch size for outer loop | 200 | {200} |
| $\lambda_f$ | Spike rate regularization | 0.25 | [0.1 ,1] |
| $f_{\text{target}}$ | Target firing rate | 20 Hz | {20 Hz} |
| $f_{\text{target}}$ for LSG | Target firing rate for LSG | 10 Hz | {10 Hz, 20 Hz} |
| $v_{\text{th}}$ | Firing threshold | 0.4 | {0.4} |
| $\lambda_v$ | Voltage range regularization | 0 | {0} |
| $N_{\text{LN}}$ | Number of neurons in the LN | 400 | {400} |

first selected a target class, and sampled two images without replacement. One of these images was to be presented in phase 1 and the remaining second image was to appear in phase 2 at a random position. Four other images were sampled randomly from the remaining classes and used to fill up the sequence in phase 2.

Classes were provided by the Omniglot[1] data set, which consists of a total of 1623 classes and a total of 32460 images (20 per class). We split up the data set according to the splits proposed in Tensorflow Datasets[2] into 964 training classes and 659 testing classes.

To extract features from the high dimensional visual input, we used a convolutional neural network (CNN) consisting of McCulloch-Pitts neurons. We defined output of a McCulloch-Pitts neuron by:

$$z = H(c - 1.2), \tag{S18}$$

where $z$ is the output of the neuron, $H$ is the Heaviside step function and $c$ is the weighted sum of inputs. The variable $c$ (weighted sum of inputs) was different for each neuron in the CNN and was given through convolutional weight structures. Our CNN neurons can be viewed to have a threshold of 1.2.

The CNN was organized into three layers with 16, 32 and 64 filters respectively, resulting in a total of 15488 McCulloch-Pitts neurons. The kernel size used for the convolutional filters was 3 by 3. Convolutions used a valid padding scheme and strides of 1. Average pooling layers (with a window size of 2) and batch normalization layers were also used to improve optimization in the outer loop. Note that both the average pooling layers and batch normalization layers can be absorbed into the CNN weights after the outer loop training. We refer to Tab. S2 for a summary of the organization of the CNN.

**Architecture** The LN consisted of $N_{\text{LN}}$ neurons and the LSG consisted of $N_{\text{LSG}}$ neurons. A fraction of $q_{\text{adapt frac}}$ of the neurons was chosen to be ALIF. See Tab. S3 for the values used in each experiment.

**Input encoding** The input $\boldsymbol{x}^t$ to the LN consisted of the output of the CNN. In addition, $\boldsymbol{x}^t$ included the phase ID encoded as a binary value, which assumed 0 in phase 1 and a value of 1 for phase 2. As the output of the CNN was 576 dimensional, the input $\boldsymbol{x}^t$ had a total of 577 dimensions.

[1]https://github.com/brendenlake/omniglot/

[2]https://www.tensorflow.org/datasets

Table S2: Architecture summary of the CNN.

| | |
|---|---|
| Weight structure | Convolution, 16 filters |
| | Batch normalization |
| 26x26x16 McCulloch-Pitts neurons | |
| Weight structure | Convolution, 32 filters |
| | Average pooling |
| | Batch normalization |
| 11x11x32 McCulloch-Pitts neurons | |
| Weight structure | Convolution, 64 filters |
| | Average pooling |
| | Batch normalization |
| 3x3x64 McCulloch-Pitts neurons | |

Images were presented for consecutive $t_{\mathrm{img}}$ time steps (20 ms in our experiments) to the CNN. Thus $\boldsymbol{x}^t$ was constant for $t_{\mathrm{img}}$.

**Human Baseline** For our estimate of the human baseline, we presented every image in the sequence for 1.5 seconds to human subjects. The code for the human test is available in the code supplements, along with a readme file containing instructions on how to run the test.

***Restricted natural e-prop*** For *restricted natural e-prop*, a broadcast of the derivative of the cross entropy loss with respect to the value of the LN output was used as the learning signal. This is expressed by:

$$L_j^{20} = B_j \cdot (\sigma(y^{20}) - 1), \tag{S19}$$

where $B_j$ represents the broadcast weights to the neuron $j$ in the LN, $\sigma$ denotes the sigmoid function and $y^{20}$ denotes the output of the LN at time $t = 20$ ms, which is the last time step in which the target class in phase 1 is presented to the model. The broadcast weights $B_j$ were optimized during the outer loop training process.

**Details of outer loop optimization** To improve the the outer loop training procedure, we applied image preprocessing and data-augmentation. We downscaled the images to a size of 28 by 28 pixels, and inverted the gray scale value. For data-augmentation, we applied three transformations to images: All images of randomly selected classes were randomly rotated by multiples of 90° and/or flipped in horizontal and vertical directions with a probability of 0.5. In addition, images were shifted randomly by up to $\pm 10\%$ in all directions. Note that rotations and flips affected all images of a class, hence effectively enlarging the available amount of training classes.

The outer loop optimized a loss function $E_C$ based on cross entropy. To define it, we introduce a label $l^n$ for every image $n$ shown in phase 2, which assumes a value of 1 only for images that belong to the same class of the image shown in phase 1 and assumes a value of zero otherwise. The loss function $E_C$ is then expressed by:

$$E_C = \sum_{n=1}^{5} -l^n \log \sigma(y^{20+20\cdot n}) - (1 - l^n) \log(1 - \sigma^{20+20\cdot n}) \,. \tag{S20}$$

Note that the output of the LN only counts after each image was fully presented.

We also optimized the weights of the CNN in the outer loop. We replaced the derivative of the Heaviside that occurs for McCulloc-Pitts neurons with the same "pseudo-derivative" as introduced in S2.1 in the backward pass.

Outer loop optimization required approximately 22 hours using a single Nvidia Tesla P100 (exact duration also depends on some hyperparameters).

**Hyperparameters** A complete list of hyperparameters that was used is given in Tab S3.

Table S3: List of hyperparameters used in application 2. Note that $N_{\text{batch}}$, $d$, $t_{\text{refrac}}$, $N_{\text{LN}}$, $N_{\text{LSG}}$ and $t_{\text{img}}$ are integer parameters.

*Natural e-prop*

| Symbol | Name | Value | Search range |
|---|---|---|---|
| $\eta$ | Learning rate of inner loop | $1.915 \cdot 10^{-3}$ | $[10^{-3}, 10^{0}]$ |
| $N_{\text{LN}}$ | Learning network size | 447 | $[100, 550]$ |
| $q_{\text{adapt frac}}$ | Fraction of neurons which use SFA | 40.5% | $[0.1, 0.5]$ |
| $\beta$ | Impact of threshold adaptation | 0.4902 | $\{0.3, 0.5\}$ |
| $N_{\text{batch}}$ | Batch size for outer loop | 285 | $[128, 512]$ |
| $N_{\text{LSG}}$ | Learning signal generator size | 239 | $[100, 500]$ |
| $\tau_{\text{learning signals}}$ | Readout-time constant learning signals | 10 ms | $\{10\,\text{ms}\}$ |
| $f_{\text{target}}$ for LSG | Target firing rate for LSG | 20 Hz | $\{20\,\text{Hz}\}$ |

*Restricted natural e-prop*

| Symbol | Name | Value | Search range |
|---|---|---|---|
| $\eta$ | Learning rate of inner loop | $9.662 \cdot 10^{-2}$ | $[10^{-3}, 10^{0}]$ |
| $N_{\text{LN}}$ | Learning network size | 456 | $[100, 550]$ |
| $q_{\text{adapt frac}}$ | Fraction of neurons which use SFA | 24.7% | $[0.1, 0.5]$ |
| $\beta$ | Beta | 0.415 | $[0.3, 0.5]$ |
| $N_{\text{batch}}$ | Batch size for outer loop | 178 | $[128, 512]$ |

Common parameters

| Symbol | Name | Value | Search range |
|---|---|---|---|
| $\tau_m$ | Membrane time constant | 15 ms | $\{15\,\text{ms}\}$ |
| $\tau_{\text{out}}$ | Readout-time constant | 10 ms | $\{10\,\text{ms}\}$ |
| $d$ | Synaptic transmission delay | 1 ms | $\{1\,\text{ms}\}$ |
| $t_{\text{refrac}}$ | Duration of refractory period | 5 ms | $\{5\,\text{ms}\}$ |
| $f_{\text{target}}$ | Target firing rate | 20 Hz | $\{20\,\text{Hz}\}$ |
| $\eta_{\text{outer loop}}$ | Learning rate of outer loop | $2 \cdot 10^{-3}$ | $\{2 \cdot 10^{-3}\}$ |
| $\lambda_f$ | Spike rate regularization | 1.0 | $\{1.0\}$ |
| $v_{\text{th}}$ | Threshold | 1.0 | $\{1.0\}$ |
| $\lambda_v$ | Voltage regularization | $10^{-2}$ | $\{10^{-2}\}$ |
| $t_{\text{img}}$ | Number of time steps per image | 20 ms | $[20\,\text{ms}, 50\,\text{ms}]$ |
| $\tau_{\text{a}}$ | Adaptation time constant | 200 ms | $\{200\,\text{ms}\}$ |

#### S6 Details of application 3: learning of posterior probabilities

**Details of the task** Each task $C$ consisted of 3 signal sources (Gaussian processes) with time-varying mean. The time-varying mean of a signal source was itself generated as a Gaussian process. We used the same type of kernel function $k(x, x'; \sigma, l, s)$ to generate the covariance matrix both for the mean of the signal source, and for generating samples of a given signal source. $k$ was given by a rational quadratic kernel:

$$k(x, x'; \sigma, l, s) = \sigma^2 \left(1 + \frac{(x - x')^2}{2sl^2}\right)^{-s}, \tag{S21}$$

and was evaluated at pairs of 5 time points $\{0\text{ ms}, 40\text{ ms}, \ldots, 200\text{ ms}\}$. The resulting covariance matrix for a 5 dimensional Gaussian distribution was then conditioned to be 0 in the first and last component, yielding a 3x3 covariance matrix $\Sigma(\sigma, l, s)$. For this task, we used $s = 0.3$. The time-varying mean $\boldsymbol{\mu}_i$ of the signal source $S_i$ was sampled in each task $C$ according to:

$$\boldsymbol{\mu}_i \sim \mathcal{N}\Big(0, \Sigma(\sigma_{\text{task}}, l_{\text{task}}, s)\Big), \tag{S22}$$

with $\mathcal{N}$ denoting a Gaussian distribution and $\sigma_{\text{task}}$, $l_{\text{task}}$ denoting the parameters used for generating tasks. In order to generate signals, we first randomly select a signal source $i$ with uniform probability among the 3 available. Subsequently, we generate a sample $\boldsymbol{w} = (w_1, w_2, w_3)$ from a Gaussian distribution with mean $\boldsymbol{\mu}_i \in \mathbb{R}^3$ and covariance matrix $\Sigma(\sigma_{\text{sample}}, l_{\text{sample}}, s)$. Note that signal samples have different parameters $\sigma_{\text{sample}}, l_{\text{sample}}$ for their covariance matrix. Finally, the obtained

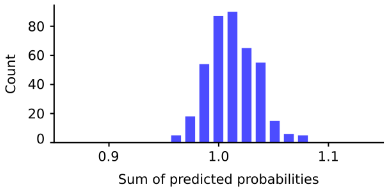

Figure S1: Histogram of the sum of LN predictions (400 samples).

sample $\boldsymbol{w}$ defines the signal $U$. Thus, signal samples were generated according to:

$$\boldsymbol{w} \sim \mathcal{N}\Big(\boldsymbol{\mu}_i, \Sigma(\sigma_{\text{sample}}, l_{\text{sample}}, s)\Big) \quad \text{with } i \sim \text{Uniform}(\{1,2,3\}) \ , \tag{S23}$$

$$U = (\underbrace{0,\ldots,0}_{\times 40}, \underbrace{w_1,\ldots,w_1}_{\times 40}, \underbrace{w_2,\ldots,w_2}_{\times 40}, \underbrace{w_3,\ldots,w_3}_{\times 40}, \underbrace{0,\ldots,0}_{\times 40}) \ . \tag{S24}$$

**Architecture** The LN consisted of $N_{\text{LN}}$ neurons and the LSG consisted of $N_{\text{LSG}}$ neurons. In both RSNNs a fraction of $q_{\text{adapt frac}}$ neurons were chosen to be ALIF. See Tab. S4 for the values used in each experiment.

**Input encoding** The input $\boldsymbol{x}^t$ consisted of a total of 101 components at time step $t$. The first 100 components of $\boldsymbol{x}^t$ were used to encode the current scalar value of the signal $u^t$, and the last component switched from 0 to 1 after 120 ms. More precisely, the input $\boldsymbol{x}^t$ was defined by:

$$x_i^t = \sin\left(\frac{u^t}{10^{2(i-1)/100}}\right) \qquad \text{for } i \in \{1,\ldots,50\}, \tag{S25}$$

$$x_i^t = \cos\left(\frac{u^t}{10^{2(i-51)/100}}\right) \qquad \text{for } i \in \{51,\ldots,100\} \ , \tag{S26}$$

$$x_{101}^t = H(t - 120 \text{ ms}) \ . \tag{S27}$$

***Restricted natural e-prop*** The learning signals in *restricted natural e-prop* were only nonzero in the last time step of processing signals $U$ (at $t = 200$ ms). For neuron $j$ the learning signal for that last time step was defined by:

$$L_j^{200} = \sum_{k=1}^{3} B_{jk}(y_k^{200} - l_k) \ . \tag{S28}$$

Here, $y_k^{200}$ denotes the final prediction of the LN and $l_k$ is 1 if $k$ is the true source of the signal and is 0 otherwise. Note that learning signals were only provided for the 45 supervised signal samples during learning a task $C$. During testing on 5 more samples in the same task, no learning signals were given.

**Details of outer loop optimization** The LN started from the same initial state when processing different signals $U$. This initial state was defined by $v_j^0 = 0$ and $a_j^0 = 0$ for each neuron $j$ in the LN. The LSG similarly started from the same initial state, where all neurons in the LSG satisfied $v_j^0 = 0$ and $a_j^0 = 0$.

In order to describe the loss $E_C$ that was optimized in the outer loop we consider the 5 samples on which the LN was tested, after the weight update had been applied in the inner loop. Let the final prediction of the LN for the $n_{\text{test}}$-th testing sample be denoted by $y_k^{(n_{\text{test}},\ 200)}$. Similarly, let the true posterior probabilities for the $n_{\text{test}}$-th testing sample be denoted by $p_k^{n_{\text{test}}}$. The loss $E_C$ was then defined by the cross entropy between the true posterior probabilities and the normalized prediction of the LN, which is defined as $\frac{y_k^{(n_{\text{test}},\ 200)}}{\sum_m y_m^{(n_{\text{test}},\ 200)}}$. Additionally, we introduced a constraint that the LN

Table S4: List of hyperparameters used in application 3. Note that $N_{\text{batch}}$, $d$, $t_{\text{refrac}}$, $N_{\text{LN}}$ and $N_{\text{LSG}}$ are integer parameters.

*Natural e-prop*

| Symbol | Name | Value | Search range |
|---|---|---|---|
| $\tau_{\text{out}}$ | Readout-time constant | 34.87 ms | [20 ms, 40 ms] |
| $\tau_{\text{learning signals}}$ | Readout-time constant of learning signals | 19.64 ms | [5 ms, 20 ms] |
| $\eta_{\text{outer loop}}$ | Learning rate of outer loop | $7.17 \cdot 10^{-4}$ | $[10^{-4}, 10^{-3}]$ |
| $\eta$ | Learning rate of inner loop | $8.36 \cdot 10^{-5}$ | $[8 \cdot 10^{-5}, 10^{-2}]$ |
| $q_{\text{adapt frac}}$ | Fraction of neurons which use SFA | 54.2% | [0.4, 1] |
| $\beta$ | Impact of threshold adaptation | 0.322 | [0.25, 0.35] |
| $\tau_a$ | Time constant of threshold adaptation | 173.9 ms | [160 ms, 190 ms] |
| $\lambda_f$ | Firing rate regularization | 0.168 | [0.01, 0.5] |
| $\lambda_v$ | Voltage range regularization | 0.0341 | [0.01, 0.5] |
| $N_{\text{LN}}$ | Number of neurons in the LN | 400 | {400} |
| $N_{\text{LSG}}$ | Number of neurons in the LSG | 300 | {300} |

*Restricted natural e-prop*

| Symbol | Name | Value | Search range |
|---|---|---|---|
| $\tau_{\text{out}}$ | Readout-time constant | 31.19 ms | [20 ms, 40 ms] |
| $\eta_{\text{outer loop}}$ | Learning rate of outer loop | $7.04 \cdot 10^{-4}$ | $[10^{-4}, 10^{-3}]$ |
| $\eta$ | Learning rate of inner loop | $1.84 \cdot 10^{-4}$ | $[8 \cdot 10^{-5}, 10^{-2}]$ |
| $q_{\text{adapt frac}}$ | Fraction of neurons which use SFA | 57.1% | [0.4, 1] |
| $\beta$ | Impact of threshold adaptation | 0.321 | [0.25, 0.35] |
| $\tau_a$ | Time constant of threshold adaptation | 178.1 ms | [160 ms, 190 ms] |
| $\lambda_f$ | Firing rate regularization | 0.233 | [0.01, 0.5] |
| $\lambda_v$ | Voltage regularization | 0.1072 | [0.01, 0.5] |
| $N_{\text{LN}}$ | Number of neurons in the LN | 400 | {400, 500} |

Common hyperparameters

| Symbol | Name | Value | Search range |
|---|---|---|---|
| $N_{\text{batch}}$ | Batch size for outer loop | 12 | {12} |
| $\tau_m$ | Membrane time constant | 15 ms | {15 ms} |
| $v_{\text{th}}$ | Base firing threshold | 1 | {1} |
| $d$ | Synaptic transmission delay | 2 ms | {2 ms, 5 ms} |
| $t_{\text{refrac}}$ | Duration of refractory period | 2 ms | {2 ms, 5 ms} |
| $f_{\text{target}}$ | Target firing rate | 20 Hz | {20 Hz} |
| $f_{\text{target}}$ for LSG | Target firing rate for LSG | 20 Hz | {20 Hz} |
| $\sigma_{\text{task}}$ | Variability of signal means | 0.5 | - |
| $\sigma_{\text{sample}}$ | Variability of signal samples | 0.6 | - |
| $l_{\text{task}}$ | Length scale of signal means | 70 ms | - |
| $l_{\text{sample}}$ | Length scale of signal samples | 40 ms | - |
| $s$ | Scale mixture for kernel | 0.3 | - |

produces normalized probability outputs. Together this yields the representation of the loss function $E_C$ as:

$$E_C = \sum_{n_{\text{test}}=1}^{5} \left[ -\sum_{k=1}^{3} p_k^{n_{\text{test}}} \log y_k^{(n_{\text{test}},\, 200)} + \log \left( \sum_{k=1}^{3} y_k^{(n_{\text{test}},\, 200)} \right) \right] + E_{\text{norm}} \,, \tag{S29}$$

$$E_{\text{norm}} = \sum_{n_{\text{test}}=1}^{5} \left( \sum_{k=1}^{3} y_k^{(n_{\text{test}},\, 200)} - 1 \right)^2 \,. \tag{S30}$$

See Fig. S1 for a histogram of the resulting normalization $\sum_{k=1}^{3} y_k^{200}$.

The outer loop optimization required approximately 72 hours for a total of $30 \cdot 10^3$ iterations on a single Nvidia Tesla P100.

**Hyperparameters** Hyperparameters were tuned separately but with the same computing budged for *natural e-prop* and *restricted natural e-prop* according to random search. We list the resulting hyperparameters in Tab. S4.
